# A simple method for methanol quantification in Spirits using UV Visible Spectroscopy and FTIR

**DOI:** 10.1101/2024.08.12.607685

**Authors:** Ronick Spenly Shadrack, Krishna Kumar Kotra, Daniel Tari, Hancy Tabi, Jacinta Botleng, Rolina Kelep, Ladyshia Regenvanu

**Author notes:** Corresponding author: Ronick Spenly Shadrack.

## Abstract

Although standards methods of food safety assessment are important, these methods are expensive and requires intensive work and time. Quality assessment for high alcohol in spirits is still a challenged for industries in developing states due to lack of financial support and technical assistance. Ultra violet visible spectroscopy (UV VIS) and Fourier Transform Infrared Spectroscopy (FTIR) offers the low cost alternative testing methods that are affordable with a short turnaround time for dissemination of results. In this work, methanol content in ethanol was assessed in two approaches using UV VIS and FTIR spectroscopy. For UV VIS method, Potassium dichromate was used as the chromogenic reagent. In FTIR, calibration curve was built by increasing methanol ration from 0 to 40% (m V^-1^) at the expense of ethanol while keeping deionised water (DO) constant at 5% (m V^-1^) concentration. This helps gauge the change in methanol concentration relative to ethanol. Results of analysis using UV VIS showed a strong negative correlation for Methanol concentration and absorbance value at UV region from 900 to 1100 cm^-1^ (r = 98.00, RMSE = 0.023) relative to increasing ethanol concentration. A strong peak was observed for methanol concentration at spectral region of 970 cm ^-1^ which is related to the methanoic acid C-O bond. The FTIR spectra region at 900 to 1050 cm ^-1^ was used for observing methanol concentration with absorbance. A strong correlation was established from spectral region of 1010 cm-1 to 1026 cm-1, enabling quantification of methanol (r= 0.99, RMSEC = 0.55). Methanol peak was observed at 1020 cm^-1^ region of the spectrum. A set of experimental repetition was made to determine limit of detection (LOD) for UV VIS and FTIR methods which was observed at 0.29 and 0.5 % (m V^-1^), respectively. The limit of quantification was 0.89 and 1.5 % (m V^-1^) for UV VIS and FTIR respectively. This study has reaffirmed the utilisation of UV VIS and FTIR as considerable alternative method for quality control of high alcohol in distilled spirits.

## Introduction

Infrared spectroscopy and UV Vis are two well established tools in analytical chemistry. Both instrument offers non-invasive and non-destructive method of analysis that is rapid and applicable to wide range of sample types (1, 2). Historically, UV VIS and IR spectroscopy have been utilised extensively due to low price of instruments, low operating cost and short turnaround time for results. Although IR spectroscopy has some advantages over UV VIS in terms of wide range of sample types, they are both important in food industry for quality control measures. Some challenges with IR spectroscopy includes interpretation of spectral data from complex mixtures, and the need to develop and maintain robust quantitative calibration model (3). The use of UV VIS and FTIR Spectroscopy in quantification of higher alcohol is often neglected due to low performance in terms of accuracy, limit of detection (LOD), limit of quantification (LOQ) and shifting region of interferogram due to thermal and structural properties of alcohol mixtures compared to recognised Gas chromatography mass spectrometry (GC-MS) method. These challenges prompt the need to study the use of UV VIS and FTIR in quantification of higher alcohol in distilled spirits considering the change of methods. The use of FTIR and UV VIS in developing countries still remains high in the food monitoring sector as most countries can afford the operating cost (4). This study investigated the use of potassium dichromate as indicator for quantification of methanol in distilled spirit using UV VIS method. The use of potassium dichromate as chromatographic indicator for calibration of UV VIS and the quantification of methanol in Biodiesel washing wastewater have been reported elsewhere in the literature (5, 6). On the other hand, the quantification of methanol in distilled spirit with FTIR is understudied and results are incomparable. The current study aims to investigate a design used in FTIR to develop calibration model that address the shifting and overlap alcohol spectral region due to structural and thermal properties of alcohol mixtures.

In the Pacific island countries including Vanuatu, distilled spirit market is relatively new and the need for advance technology with higher operating cost such as GC-MS for quantification of methanol in spirits is not practical, though necessary. It has been confirmed that alcohol is a leading risk factor for disease and injury in the Pacific island countries (7, 8). Thus, it is just important to investigate simple and rapid method for high alcohol quantification such as methanol in distilled spirit to meet safe limit recommended by international food regulatory organisation including international standards organisation (ISO), Food and Drugs Association of America (FDA) and European food association. In the European countries the limit for methanol in distilled spirit is 10 g methanol/l ethanol (9) while in the US, methanol limit in alcohol is 7 g methanol/ l ethanol (10). This study relies on two mentioned standards for quality control of methanol in distilled spirits irrespective of accurate estimation of the amount that are well below instrumental LOD. The precision and accuracy of FTIR spectra scan at every two nanometres will also be investigated for its use in quality control of higher alcohol in distilled beverages.

## Methods

### Materials

Methanol (CH_3_OH, Merck, HPLC, ≥ 99.9%), sulfuric acid (H_2_SO_4_, Merck, 97%), potassium dichromate (K_2_Cr_2_O_7_, Cinética, 99%), ethanol (CH_3_CH_2_OH, Qhemis, 99.5%), and DO water. All reagents were used as received. The distilled spirits tested in this study was donated by the 83 island distillery in Port Vila, Vanuatu. The tested sample were distilled from fermented sugarcane and were at Heads category of purification for high alcohol.

### Detection and quantification of Methanol by FTIR

Prior to collection of the calibration spectra, room environment was controlled with temperature and humidity being 21 °C and 35%, respectively. For the methanol FTIR calibration model, DO water volume was held constant at 60% (m V^-1^) of the calibration solution while methanol concentration increases from 1.8 to 40% (m V^-1^) at the expense of ethanol in 21 data points. The blank was made of deionised water only. The mid IR spectra by Fourier transform were obtained by a Bruker spectrometer, model Alpha II, coupled with ATR single-reflectance cell, and diamond prism with a contact diameter of 1.8 mm. The FTIR instrument used was pre-calibrated to obtained spectra point at every two nanometres which equate to half of the total data for FTIR instrument that obtained data in every nanometre. Spectra were obtained in the spectral region between 500 cm^-1^ to 4000 cm^-1^ with 23 scans in absorbance mode, resolution of 4 cm^-1^, apodization by the Sqr Triangle function, and 20 μl of each sample entry was used. These operating conditions led to 20 s analysis time. Area under peaks were used as analytical signal to construct the calibration model.

### Detection and quantification of Methanol by UV VIS

The absorbance read at 900 cm^-1^ to 1100 cm^-1^ were obtained using diffuse reflectance spectroscopy in the visible region (325-1100 cm^-1^) in a Shimadzu Spectrophotometer, model UV – 1800. To construct the standard curve, standard preparation was described as follows; methanol volume was held constant at 5% (m V^-1^) of the standard solution while ethanol concentration increases from 5 to 40% (m V^-1^) at the expense of DO in 16 data points. Before measurement in the spectrophotometer, the standards for constructing the calibration curve was prepared using the following steps; 5 ml of H_2_SO_4_ was poured into each diluted samples followed by addition of 1 ml of K_2_Cr_2_O_7_ solution (10% m V^-1^), and, finally, the solutions were mixed for approximately 2 min by vortexing. The finished mixtures were analysed using a blank sample prepared in the same way described above by pouring H_2_SO_4_ and the K_2_Cr_2_O_7_ solution into 5 mL of D0 without adding methanol.

### Statistical analysis model

Based on the FTIR and UV VIS spectra, the analysis were performed at frequency intensities recorded in the domain 1010-1026 cm ^-1^ and 900 -1100 cm ^-1^, respectively. Principle component analysis (PCA) was performed by Unscrambler 10.4 Software, version 10.4 (Camo Software AS, Oslo, Norway) for sample variability and discrimination. The peak areas from the specified wave numbers were analysed for their concentration and prediction using Partial least squares (PLS) regression with the same software. For the model development, 70 % of sample data was used in learning set and the remaining 30 % for the validation set. Spectra were cantered prior to further analysis. The PLS model was pre-weighted and a full cross validation was performed. A general linear regression model (GLM) was used to evaluate the limit of detection (LOD) and limit of quantification (LOQ) for the predictive model.

## Results and Discussion

### Identification of methanol and ethanol in the UV Vis spectra of pure standards and a spiked sample

In order to distinguish the colour differences in the pure methanol and ethanol standard under oxidation reaction with acid dichromate, a trail was made with the sub sample of known concentration following the method. The standard methanol (99.9%) (Fig.1, A) and ethanol (99.8%) (Fig.1, B) in the chemical mixtures and the blank (DO) (Fig.1, D) can be easily differentiated with the colour differences. After the oxidation of methanol, the samples were revealed a colour scale of golden yellow for blank sample (Fig.1, D) and a golden teal (Fig.1, A) for methanol. In the presence of ethanol, methanol presence by colour is challenging to differentiate due to presence of golden green colour produced from oxidation of ethanol by dichromate (Fig.1, C).

**Figure 1.**
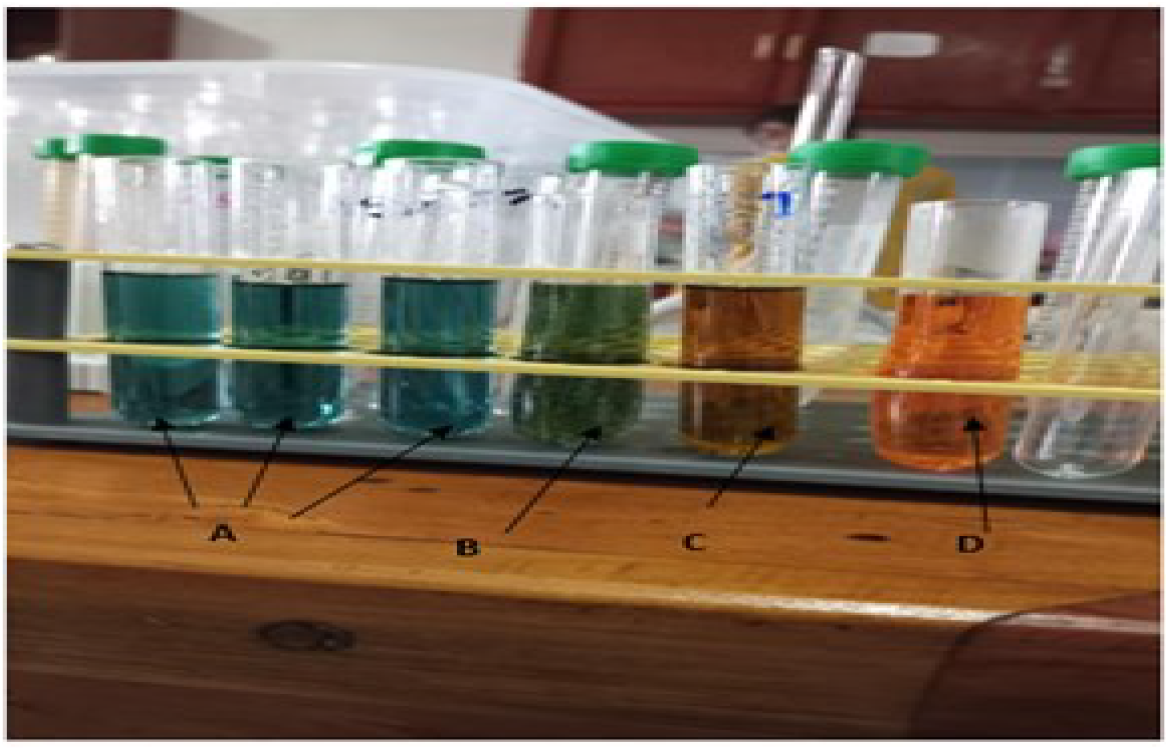
A. Methanol (99.8%) in acid dichromate, B. Ethanol (99.6%) in acid dichromate, C. Spiked methanol and ethanol (1:2 % m V-1) in DO, with acid dichromate and, D. reagent black in acid dichromate.

Collection of sample spectrum from a full spectrum scan is a common technique to determine peaks absorption of molecule in any given sample. In this experiment, in order to identify peaks relative to change in concentration of methanol and ethanol in acid dichromate, the whole sample spectrum was scanned at region of 325 cm-1 to 1100 cm-1 (Fig. 2). Ethanol change in peak intensity was observed at 600 nm while methanol change in peak intensity was strongly correlated at 970 cm-1. As the concentration of methanol decrease relative to ethanol, peak intensity reflecting the change was observed in the 970 cm-1 region of the spectrum.

**Figure 2.**
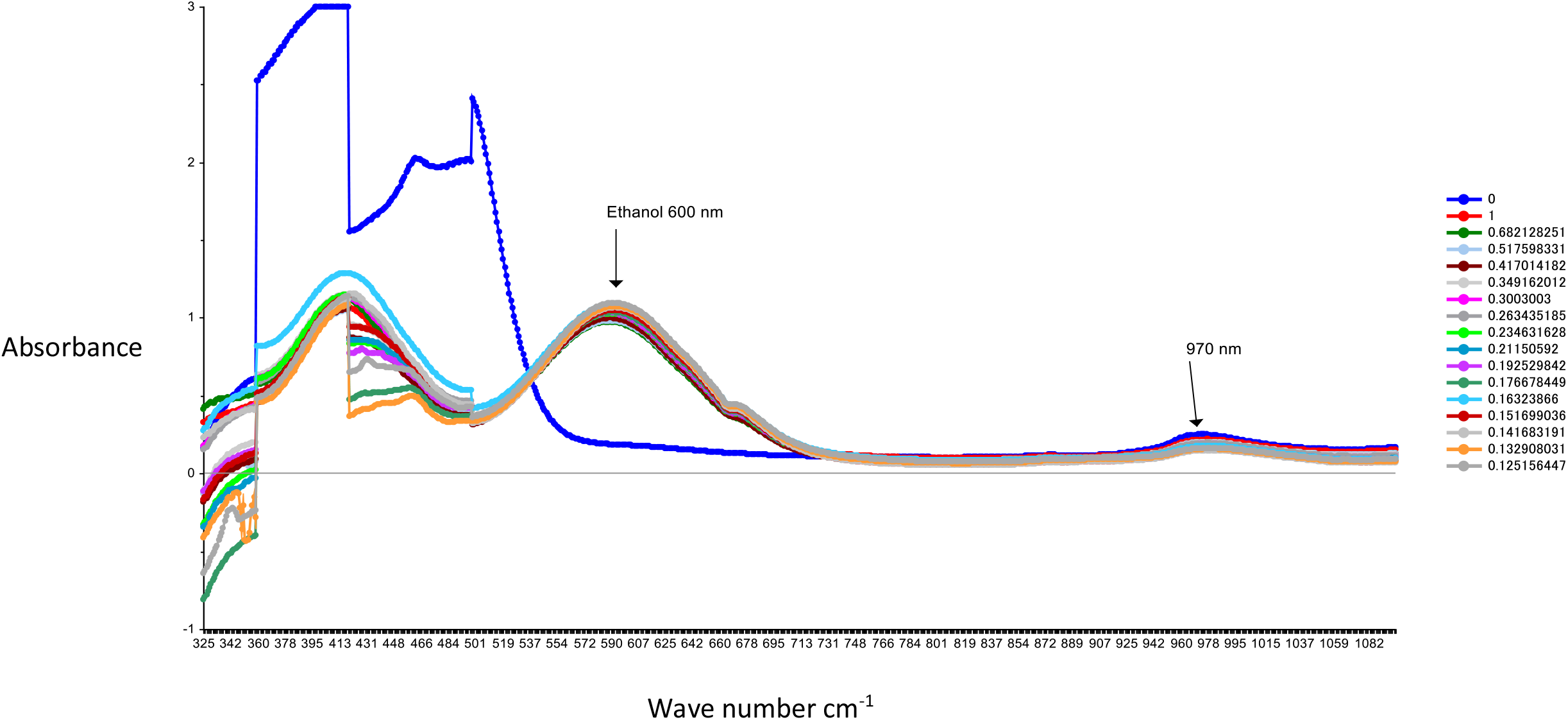
UV Vis spectra of spike methanol in Ethanol and DO using the acid dichromate method. Ethanol peaks at 600 cm-1, Methanol peaks at 970 cm-1.

After the methanol oxidation process, the samples were revealed with colours ranging from a scale of golden yellow to green (Fig1.A). As shown in Equation 1, it is suggested that methanol was wholly oxidized to CO_2_ and H_2_O in the presence of K_2_Cr_2_O_7_ under a strongly acidic medium (11) at the same time that Cr6+ (golden yellow) was reduced to Cr3+ (teal colour). It is also believed that this reduction may pass through an intermediate state of Cr4+ due to the reddish-brown coloration (11). The intensity of the colour read at 600 cm-1 was inversely proportional to the concentration of methanol in each sample observed at 970 cm-1 of the spectrum. This result is in contrast to Santos et al.,(6) who reported that methanol detection with UV Vis in biodiesel waste water was observed 600 cm-1. Here, colour intensity at 600 cm-1 was propositional to ethanol concentration.

### Calibration and validation of PLS procedure for methanol

In view of the UV Vis spectra display in fig.2, there was a gradual decrease in absorbance as the concentration of methanol decrease relative to increasing ethanol concentration. Such behaviour was reported in Yuan et al., (12)The PLS model development from the acid dichromate UV Vis method showed a good correlation with methanol concentration at 970 cm-1 spectra region with a correlation coefficient of r = 0.99 and Root mean square error coefficient (RMSEC) of 0.007 (Fig.3). The validation of the model revealed a strong correlation of r = 0.96, with RMSECV of 0.028 % (m V^-1^) (Fig.4.). This study has confirmed the UV Vis method using acid dichromate as potential chromatic agent for methanol quantification in alcohol beverages especially distilled spirits. The use of PLS model showed a good predictive model and such technique is cost effective for small businesses for quality control of methanol in alcohol beverage. The use of acid dichromate showed strong correlation with methanol concentration in waste water from Biodiesel washing observed by Santos et al.,(6).

**Figure 3.**
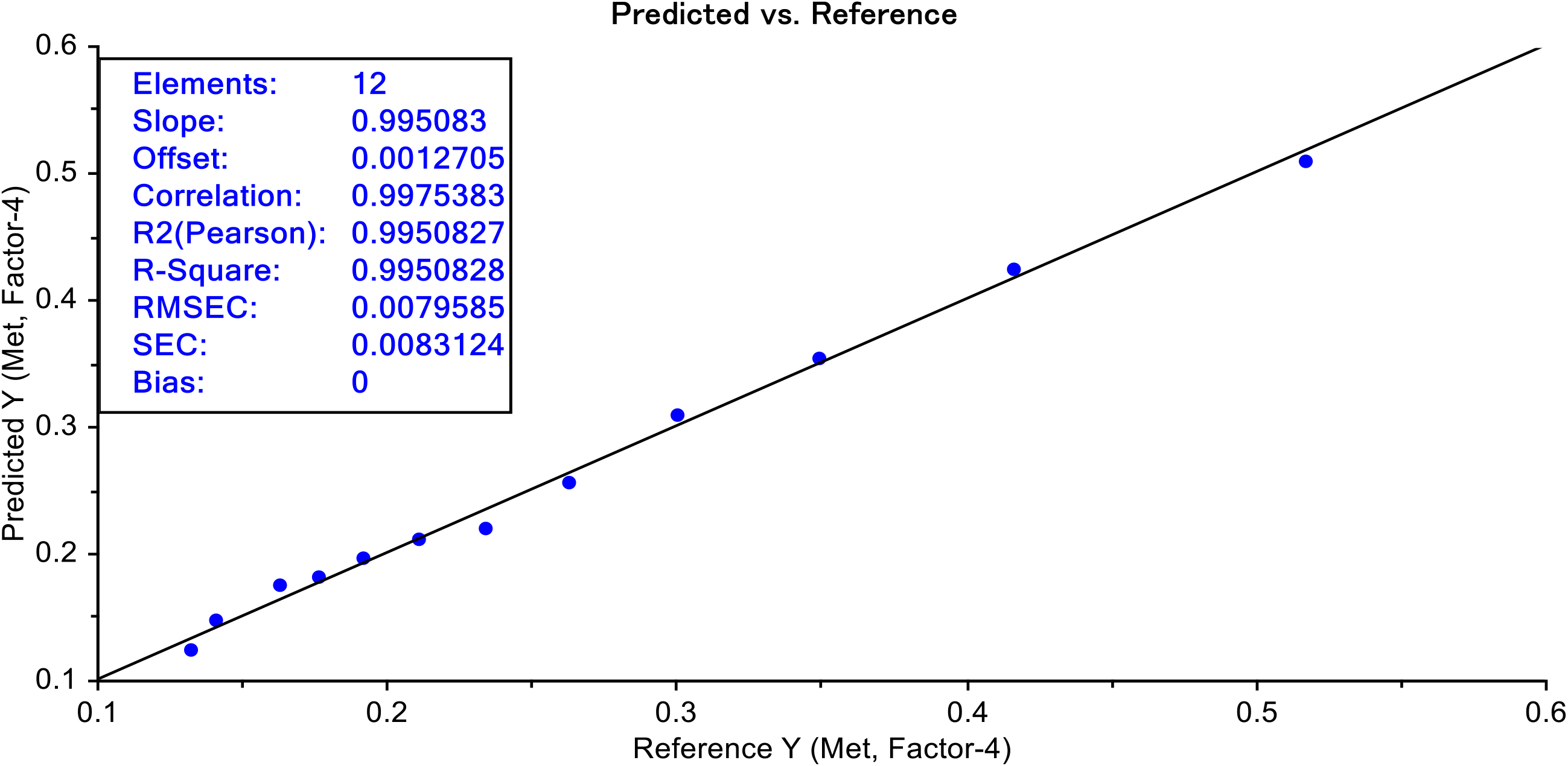
Calibration set, UV Vis (region 900-1100) using acid dichromate

**Figure 4.**
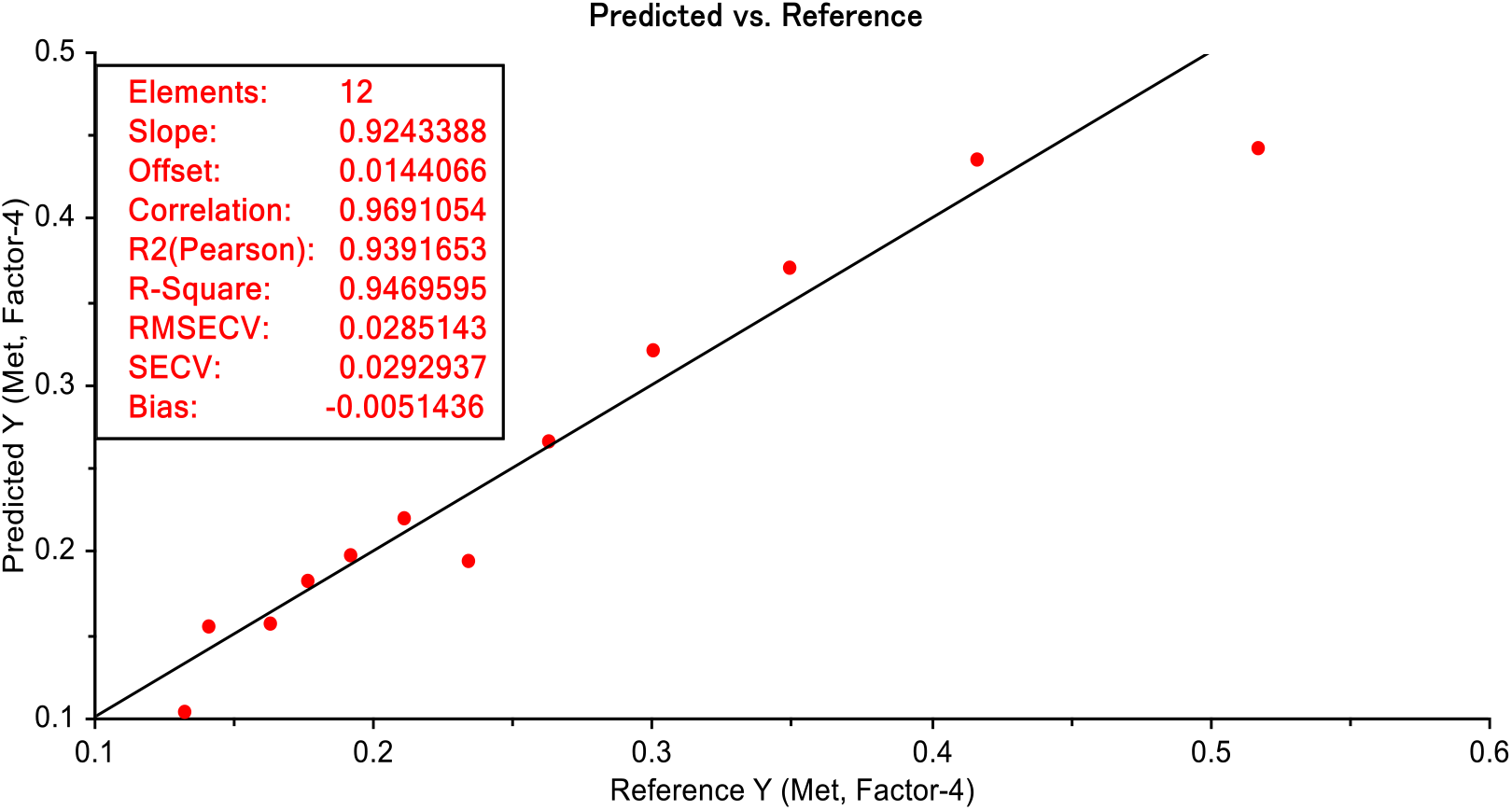
Validation set, UV Vis (Region 900-1100 nm) using acid dichromate

### Identification of methanol and ethanol in the FTIR spectra of pure standards and a spiked sample

Use of IR spectroscopy for identification of alcohol species in sample mixtures is common although the methods presented challenges with accuracy and precision compared to standard methods (3, 4, 13). The use of Near Infra-red (NIR) and Mid Infra-red spectroscopy (MIR) for quantification of ethanol in alcohol-based gel hand sanitiser during COVID 19 was possible (13), but the limit of detection (was above the required standard regulated by international organisations (9, 10). In order to build the FTIR model, spike volume of methanol at the expense of ethanol with DO held constant, the pattern recognition was developed and calibrated according to spile concentration of sample mixtures. The calibration spectra were obtained at spectra region between 1010 cm-1 to 1026 cm-1 wavelengths with frequency intensity measured in absorbance unit (Fig.5.). A strong linearity was observed for methanol peak at 1020 cm-1 used in building the calibration model. The methanol peak at 1020 cm-1 was observed for alcohol beverage using the MIR (13) and similar observation was observed in this study using FTIR spectroscopy (Figure.5.).

**Figure 5.**
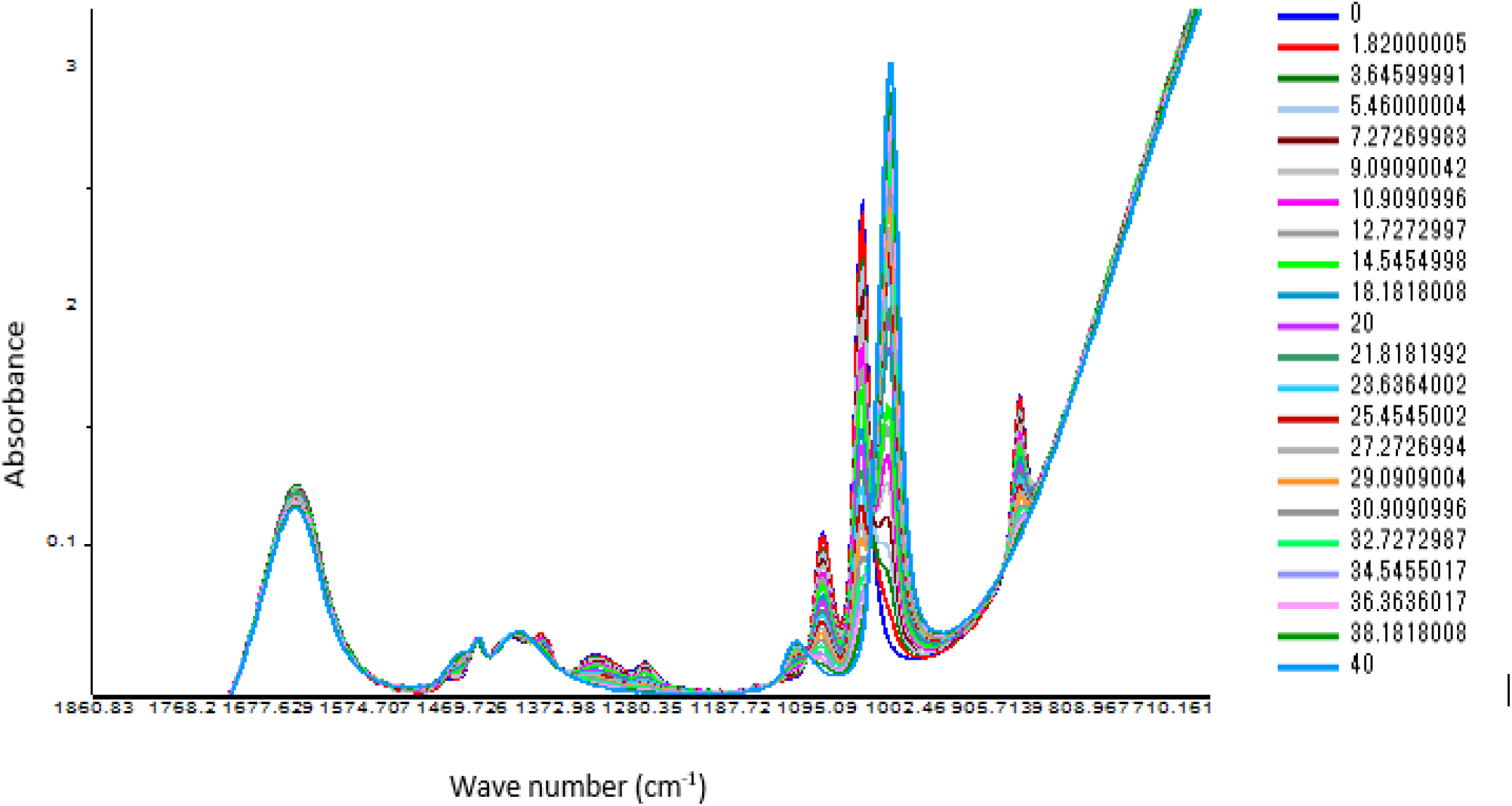
FTIR Spectral distribution for the inverse ethanol/methanol relationship. Dark blue spectra colour represents ethanol standard without methanol and light blue spectra colour represent 40% methanol.

### Calibration and validation of PLS procedure for methanol

By applying the PLS regression for methanol calibration curve (Fig. 7 and Tab. 2) we were able to predict methanol content in spiked samples and test samples. The FTIR absorbance at 1010 cm-1 to 1026 cm-1 showed a strong correlation with methanol concentration. The loading plot showed a region where methanol and ethanol concentration were equal at point 0 of the Factor 1 which explain 100% of the differences (Fig.6). Calibration model of spiked samples showed a strong linear relationship with the correlation coefficient of r = 0.997 and RMSE of 0.55% (m V^-1^). The validation model also revealed a strong linear relationship with correlation coefficient of r = 0.998 and RMSECV of 0.62% (m V^-1^) (Fig.8.). This model though was developed from a two interval spectra data point; it is still reliable for the prediction of methanol in distilled alcoholic beverages.

**Figure 6.**
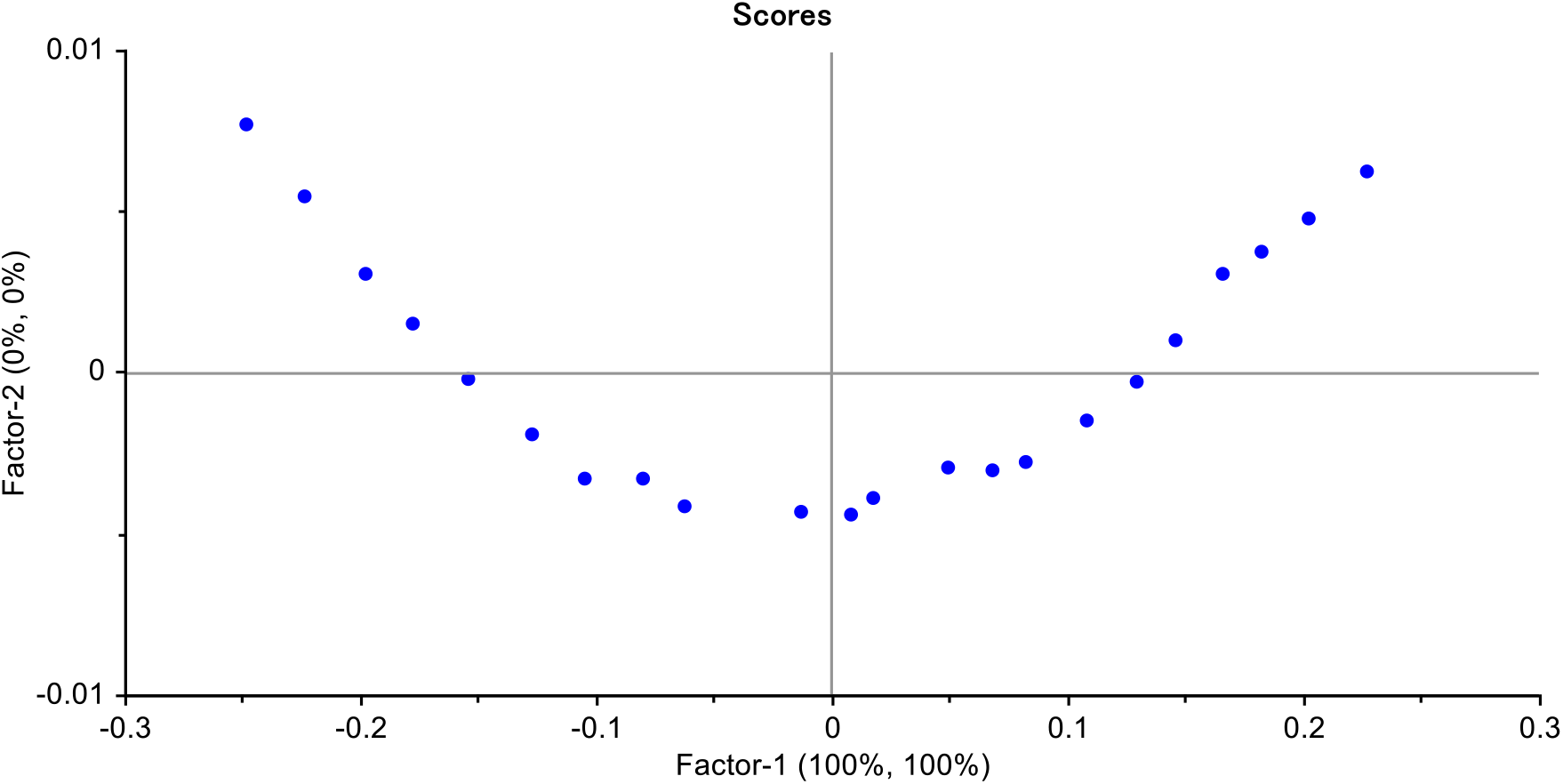
Loading plot of the Inverse relationship between ethanol and methanol used in the FTIR model development

**Figure 7.**
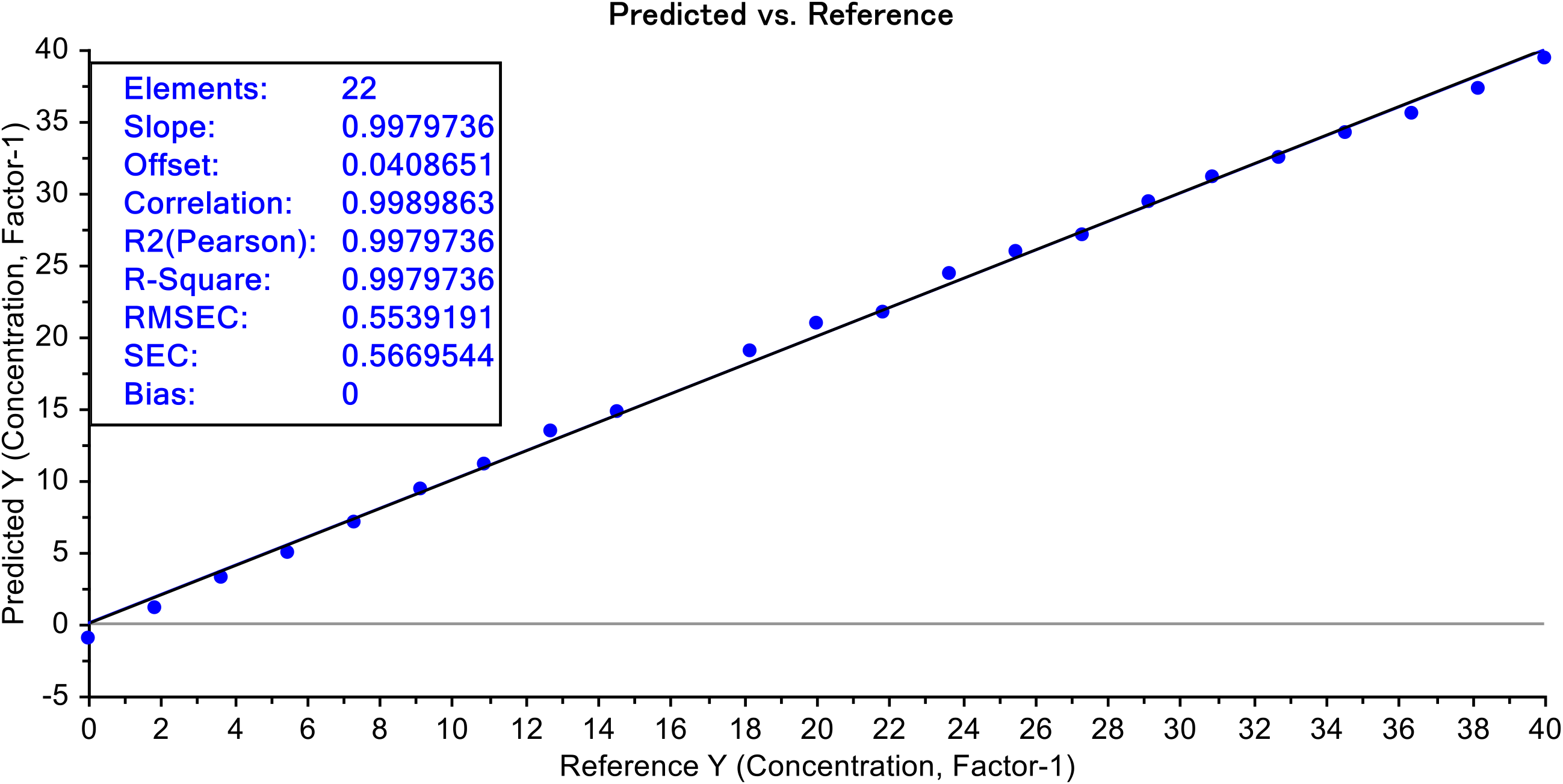
Calibration set for MET/ET using FTIR spectroscopy (1010 to 1026 cm-1)

**Figure 8.**
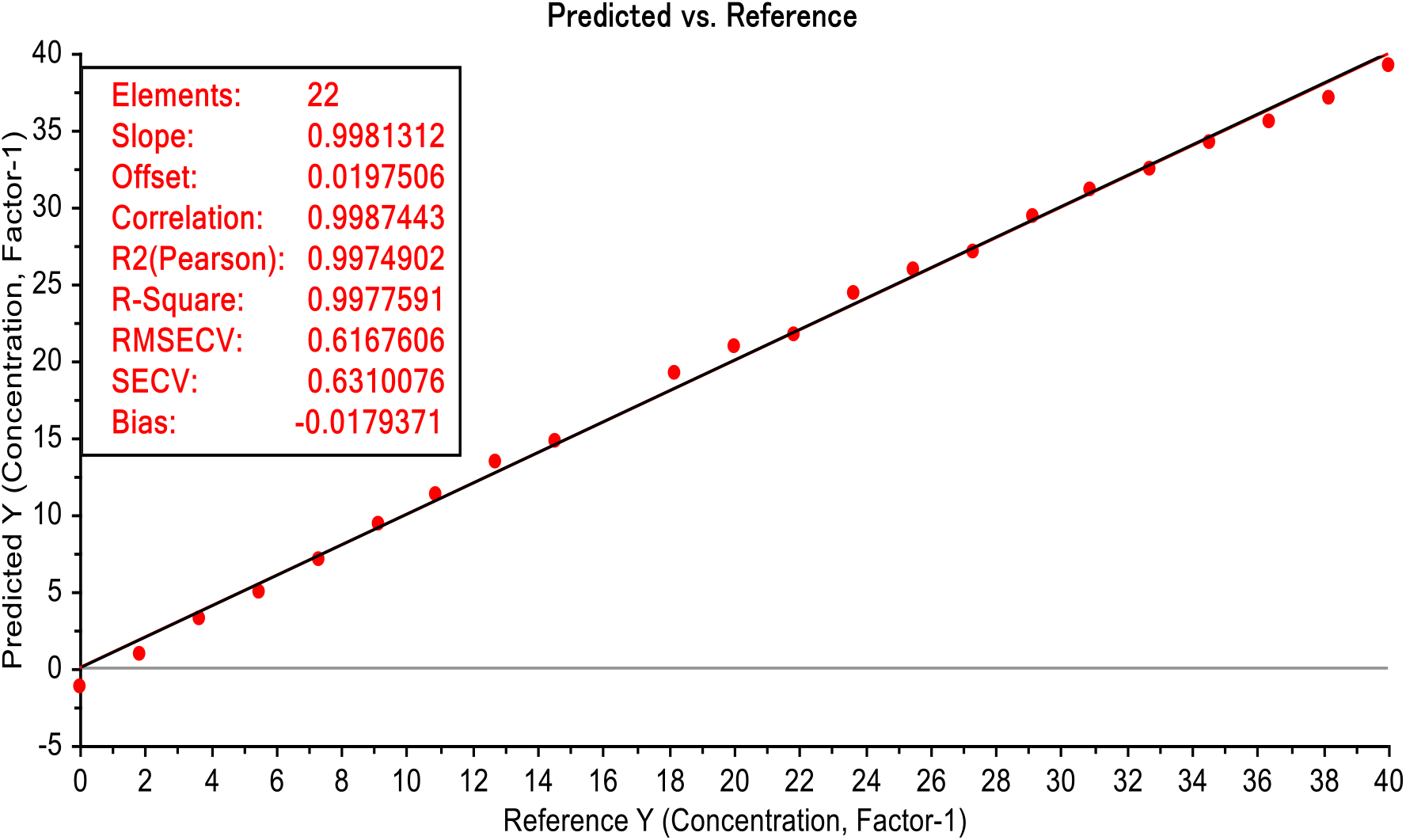
Validation curve for MET/ET using FTIR spectroscopy (1010 cm-1 to 1026 cm-1)

### 3. Analytical and statistical parameters of the analytical curve for each of the evaluated instrumental methods by Second analyst (n ≤ 16)

**Table 1.** Analytical curve for data collected using the current method.

| Instruments | Working range(%) | Analytical curve equation | r<sup>2</sup> | RSD<sub>r</sub> (%) | LOD (%) | LOQ (%) |
| --- | --- | --- | --- | --- | --- | --- |
| ATR-FTIR | 0.182-40 | $y = 0.8929x - 0.1083$ | 0.91 | 11.4 | 0.29 | 0.89 |
| UV-Vis | 0.125-1 | $y = 0.289x + 0.3271$ | 0.96 | 15 | 0.50 | 1.50 |

The analytical curve for the FTIR method showed that limit of detection was relatively low 0.29 % (m V-1) with a correlation coefficient of r = 0.91, similar to LOD reported in previous study (14). It is worth knowing that the FTIR instrument used recorded spectra data in every two nanometre making it a reliable instrument in less developed laboratory for quality control methanol in alcohol beverages. The UV Vis analytical curved from the current method showed low limit of detection at 0.5% (m V-1) making it a potential method for quality control of methanol in distilled alcohol. The spectral distribution of methanol relative to ethanol in Fig.10 revealed, most of the Heads distilled alcohol are around the same concentration with spiked methanol of 0%-1% in 40% ethanol. The spectral distribution of 100% methanol can be easily distinguished from the rest of the spiked sample and test heads samples (Fig.10). Figure 10 also showed the differences in spectral distribution around 1020 cm-1 used to build the prediction model. It is evident that quality control of methanol in distilled alcohol beverage can be easily determined using the FTIR method. A strange trend was observed in Fig.10 for spiked methanol in ethanol where the peaks was minor as spectra were almost a straight line (Met5-Eth7), reflecting the behaviour of the mixture of spiked methanol in ethanol when concentrations are at same ratio affecting the model development.

**Figure 9.**
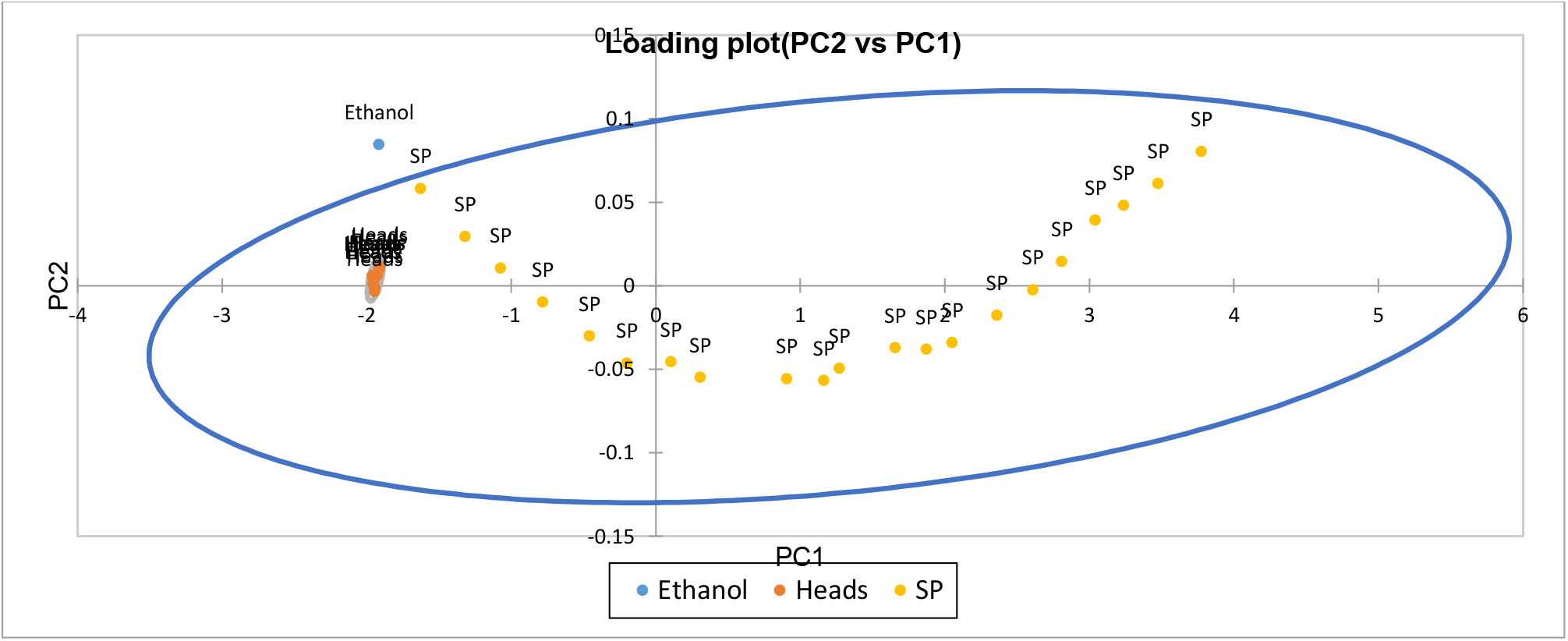
Loading plot of Principle component analysis for pure ethanol, spiked (SP) methanol and distilled alcohol (Heads).

**Figure 10.**
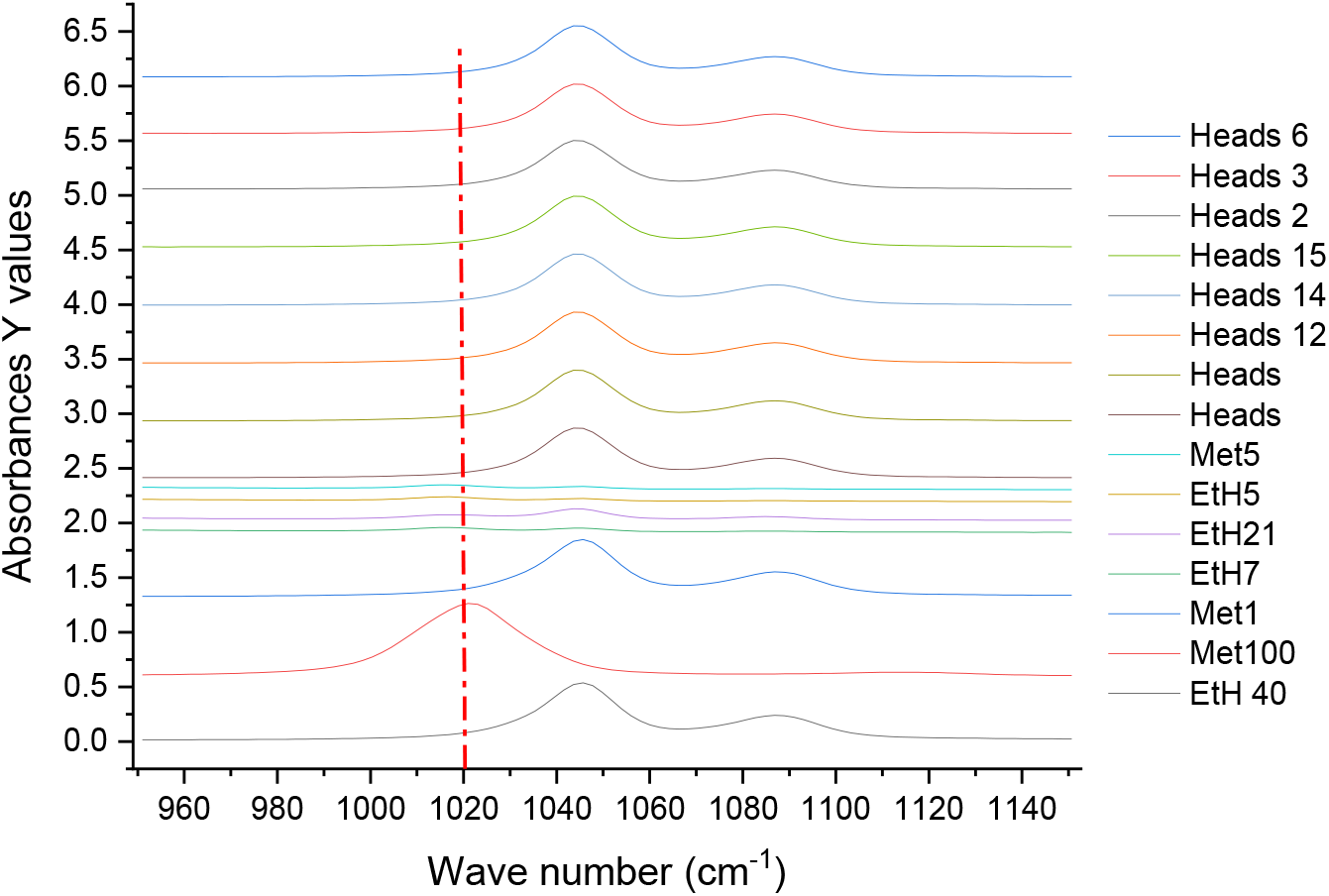
The FTIR spectral distribution of Methanol (Met.), Ethanol (EtH.) and Heads distilled alcohol beverages at 950 cm-1 to 1150 cm-1. The vertical red line (1020 cm-1) indicates the active peak for Methanol.

Due to lower concentration below the LOD of the FTIR method, differentiation of various spiked methanol in ethanol, distilled alcohol (Heads) and pure ethanol is shown in figure 9, for qualitative analysis. It was evident that the distilled alcohol (Heads) tested has methanol concentration as the distribution of samples lie within the 95% confidence eclipse, however, the amount is quite low. The data generated from the prediction model revealed negative concentration and were not presented in this study, but a qualitative test using the PCA is still adequate.

The pure methanol solution has the characteristic vibration frequency at 1020 cm-1 (major signal) while ethanol had the characteristic frequency at 1047 cm-1 (major signal) and 1087 cm-1 (minor signal), similar to the observation of Coldea et al.,(16). The selected absorbance of major and minor signals specific for pure methanol and ethanol clearly displayed in Fig.10. These frequencies for Methanol and ethanol are specific for stretching vibrations of C-O bonds in these molecules (16).

## Conclusion

UV Vis and FTIR method can be useful not only for determining the methanol and ethanol content in spirits, but also to evaluate the ratio between these two volatile components. UV Vis and FTIR/PLS offers valuable advantages when choosing versus conventional methods such as GCMS. Their importance is justified by being a rapid, efficient and non-destructive tool for screening alcoholic beverages. Besides, quantitative PLS regression analysis with FTIR spectral data is more practical in developing countries as high end instruments are not affordable. The UV Vis acid dichromate method can be utilised also with simple linear regression for peak absorbance at 970 cm^-1^ spectra region. These two method becomes important for authenticity control based on provenience region for desired distilled alcohol beverages. This study has confirmed the utilization of the two methods in quantification and qualification of distilled alcohol beverage for high alcohol.

## Acknowledgment

We acknowledge the support from Dr. Adel B. from the CEFAS UK for providing the instruments needed for this study.

## Author Contribution

R.S.S. designed and analyzed the study and wrote and reviewed the manuscript. K.K.K. analyzed the data, wrote the main text of the manuscript and reviewed the manuscript. Similarly, J.B., D.T., H.T., L.R. and R.C. wrote the main text of the manuscript and reviewed and edited the manuscript. All the authors have read and agreed to publish the manuscript.

## Data Availability and Materials

The datasets generated and analyzed during the current study are not publicly available due to property rights protection but are available from the corresponding author upon reasonable request.

## Ethics approval

The Study was approved by the Vanuatu Bureau of Standards, Vanuatu Government.

## Consent for publication

All the authors have provided consent for publication.

